# Divergent changes in perturbation-induced brain reconfiguration following depression treatment with psilocybin and escitalopram

**DOI:** 10.64898/2026.06.22.733731

**Authors:** Paulina Clara Dagnino, Irene Acero-Pousa, Robin L. Carhart-Harris, David Erritzoe, David J. Nutt, Morten L. Kringelbach, Yonatan Sanz Perl, Gustavo Deco

## Abstract

A central challenge in neuroscience is understanding how the human brain is organised to support optimal functioning and adaptability. One approach to characterise complex brain dynamics is by artificially perturbing whole-brain models. Here, we asked whether whole-brain organisation under perturbation in major depressive disorder (MDD) changes after intervention with psilocybin and escitalopram. First, we built whole-brain models of pre- and post-treatment resting-state functional magnetic resonance imaging (fMRI) and obtained an initial generative effective connectivity (GEC) matrix for each individual. Then, we employed systematic and local artificial perturbations across intensities, re-optimised each model to create a response GEC (GECr), and assessed the extent of brain reorganisation by quantifying the brain network reconfiguration index (NRI). Our results showed that the global brain NRI increases with psilocybin and decreases with escitalopram. Across sessions and interventions, higher global NRI was related with localised perturbations in brain areas orchestrating the brain’s hierarchical dynamics. Traditional approaches complemented our investigation. Our findings suggest distinct neural changes following each treatment for MDD. The increase in brain reorganisation under perturbation following psilocybin is consistent with greater brain flexibility and changeability, whereas the decrease following escitalopram suggests more stabilised brain dynamics. Overall, perturbation-induced brain NRI may represent a useful approach for uncovering neural changes following different interventions for depression.

## Introduction

In neuroscience, a central challenge is understanding how the human brain is organised to support behaviour and ensure survival. A key feature of brain dynamics is the balance between stability and flexibility, enabling the brain to maintain function while adapting to inherent internal and external signals. This balance underpins different brain states such as depression, associated with rigidity, and psychedelics, associated with flexibility and malleability (Vohryzek et al., 2022). A way of characterising such complex brain dynamics is applying artificial perturbations to whole-brain models, which has proven effective for quantifying brain sensitivity (Breyton et al., 2025; Deco et al., 2018; Sanz Perl et al., 2022), and assessing transitions between brain states (I. Acero-Pousa et al., 2025; Deco et al., 2019; Gu et al., 2017; Sanz Perl, Fittipaldi, et al., 2023; Sanz Perl, Pallavicini, et al., 2023; Zheng et al., 2025). However, these approaches have primarily evaluated the response to perturbations, leaving unresolved how brain organisation changes in the presence of sustained perturbations.

Major depressive disorder (MDD) is a neuropsychiatric disorder characterised by anhedonia, low mood, and cognitive challenges. The importance of MDD arises from its top-ranking burden of disease worldwide (GBD, 2018) and the challenge of successful treatment given its neurobiological complexity and heterogeneity of symptoms (Holtzheimer & Mayberg, 2011). Recent studies have reported treatment-specific changes in whole-brain dynamics response to local perturbations. One investigation evaluated systematic perturbation of MDD whole-brain models with psilocybin therapy (Vohryzek et al., 2024). Results uncovered candidate regions responsible for restoring healthy brain dynamics. Another investigation measured susceptibility on resting-state functional magnetic resonance imaging (fMRI) scans of MDD patients before and after treatment with psilocybin and escitalopram (Socoro-Garrigosa et al., 2025), from a double-blind phase II randomised controlled trial (Carhart-Harris et al., 2021). They built whole-brain models and perturbed brain areas locally and separately at different noise levels. Psilocybin treatment showed increased susceptibility whereas escitalopram treatment showed decreased susceptibility. Whether these treatment differences in susceptibility are reflected in how the brain reconfigures under such perturbations remains unknown.

Here, we asked whether brain network reconfiguration following perturbation of patients with MDD changes following intervention with psilocybin and escitalopram. To do so, we introduced the brain network reconfiguration index (NRI), a measure quantifying the functional and generative capacity of brain dynamics to reorganise with a perturbation. Instead of evaluating sensitivity to stimulation, we measured the capacity of the brain network to reorganise despite a perturbation. We analysed the same cohort as in (Socoro-Garrigosa et al., 2025) and built whole-brain models. For each patient, we derived an initial generative effective connectivity (GEC) matrix from the anatomical structural connectivity (SC), which was fitted to empirical fMRI data using an optimisation procedure. Then, we perturbed the models by adding noise to each brain area separately. Inspired by the two-step modelling procedure from (Irene Acero-Pousa et al., 2025) (i.e., fitting a model to the output of a previous model), we re-optimised and fitted our perturbed models to the initial GEC, thus obtaining a response GEC (GECr). We defined the NRI as the inverse similarity (i.e., decorrelation) between the response GECr and the initial GEC, with higher NRI reflecting greater reorganisation with perturbation. Overall, this framework allowed us to assess the brain network’s capacity and underlying organisation principles of network reorganisation with perturbation.

Our perturbation-induced brain network reconfiguration index (NRI) successfully showed changes with different interventions for MDD, increasing and decreasing with psilocybin and escitalopram treatment, respectively. We move beyond model-based approaches which implement sensitivity analysis in terms of response to perturbations and provide a characterisation of brain reorganisation under sustained perturbation. Overall, our findings offer new insights into the neural effects of depression treatment with psilocybin and escitalopram.

## Materials and Methods

### Trial

The trial design and primary clinical outcomes (clinicaltrials.gov: NCT03429075) have been documented in (Carhart-Harris et al., 2021). The clinical trial took place at the National Institute for Health Research Imperial Clinical Research Facility, with sponsorship from Imperial College London. It obtained ethical approval (ID 17/LO/0389) from the NHS Research and Imperial College Joint Research and Compliance Office, and approval from the Health Research Authority and Medicines and Healthcare Products Regulatory Agency. The study was performed under a Schedule 1 Drug Licence provided by the UK Home Office. Participants did not receive any final compensation and provided written informed consent.

### Treatment protocol

The final number of participants for this analysis were n=22 (mean age= 41.9 years, s.d.= 11.0, 14 male and 8 female), and n=20 (mean age= 38.7 years, s.d. =11.0, 14 male and 6 female) for the psilocybin and escitalopram arms, respectively. Baseline resting-state fMRI with eyes-closed scans were performed for all patients. In the first dosing day (DD1), participants randomly allocated to the psilocybin arm received two doses of 25 mg of psilocybin, 3 weeks apart, and participants allocated to the escitalopram arm received 1 mg of psilocybin. In an attempt to standardize expectation prior to dosing, all individuals were informed that they would receive psilocybin, but they were not given information on dosage. In the second dosing day (DD2) - three weeks after DD1 -, patients received the same dosage as in the first dosing day without crossover in dosages between the two arms. From the day after DD1 onwards, till 6 weeks and 1 day after, patients took daily capsules corresponding to one per day in the initial three weeks and two per day afterwards. In the psilocybin arm the content was inert placebo (microcrystalline cellulose) and in the escitalopram arm the content was 10 mg of escitalopram in each capsule. Post-treatment resting-state fMRI with eyes-closed scans were performed for all patients. Details on participant recruitment, MRI paradigm and rationale are provided in the **Supplementary Information**.

### Brain parcellation

All neuroimaging data was parcellated using the DK80 parcellation (Deco, 2021) which combines the Mindboggle-modified Desikan Killian parcellation of 62 cortical brain areas (31 in each hemisphere) (Desikan et al., 2006; Klein & Tourville, 2012) with the following 18 subcortical brain areas (9 in each hemisphere): hippocampus, amygdala, subthalamic nucleus (STN), global pallidus internal segment (GPi), global pallidus external segment (GPe), putamen, caudate, nucleus accumbens (NA) and thalamus.

### Brain network reconfiguration index

In order to quantify changes in the brain network reconfiguration index (NRI) (**Figure 1A**) following either intervention (**Figure 1B**), we first built a whole-brain Hopf model of each individual. We represented the local dynamics of each brain area, in a DK80 parcellation (62 cortical and 18 subcortical areas) (Deco, 2021), with a Stuart-Landau oscillator. We created a generative effective connectivity (GEC) matrix from each model, where the strengths of existing anatomical connectivity are iteratively adapted until best fit to empirical fMRI data (**Figure 1C**) (Deco et al., 2017). We used an optimisation procedure using the thermodynamic concept of “arrow of time” to capture asymmetries in the direction of information flow (Deco, 2024; Kringelbach et al., 2024; Lynn et al., 2021). For the NRI computation, schematised in **Figure 1D**, we stimulated the models *in-silico* by increasing the Gaussian noise of the Stuart-Landau oscillator of one area, whilst maintaining the rest of the areas at 0.01. Then, we re-optimised the GEC containing the perturbation by fitting it to the initial simulated measures, creating a response GEC (GECr). This way, the GECr captures whole-brain re-organisation under a constant perturbation that attempts to preserve the initially modeled configuration. Lastly, we quantified the NRI by calculating the decorrelation between the GECr and GEC. The higher the NRI, the lower the similarity between the initial model and the response model, and thus the more reorganised brain network with a given perturbation. We stimulated separately all brain areas across intensities ranging from 0.02-0.10 in steps of 0.01. We obtained the NRI across areas and stimulation intensities for each individual. Full mathematical description of the Hopf model, its linearization, optimisation, and NRI computation are provided in the **Supplementary Information**.

**Figure 1.**
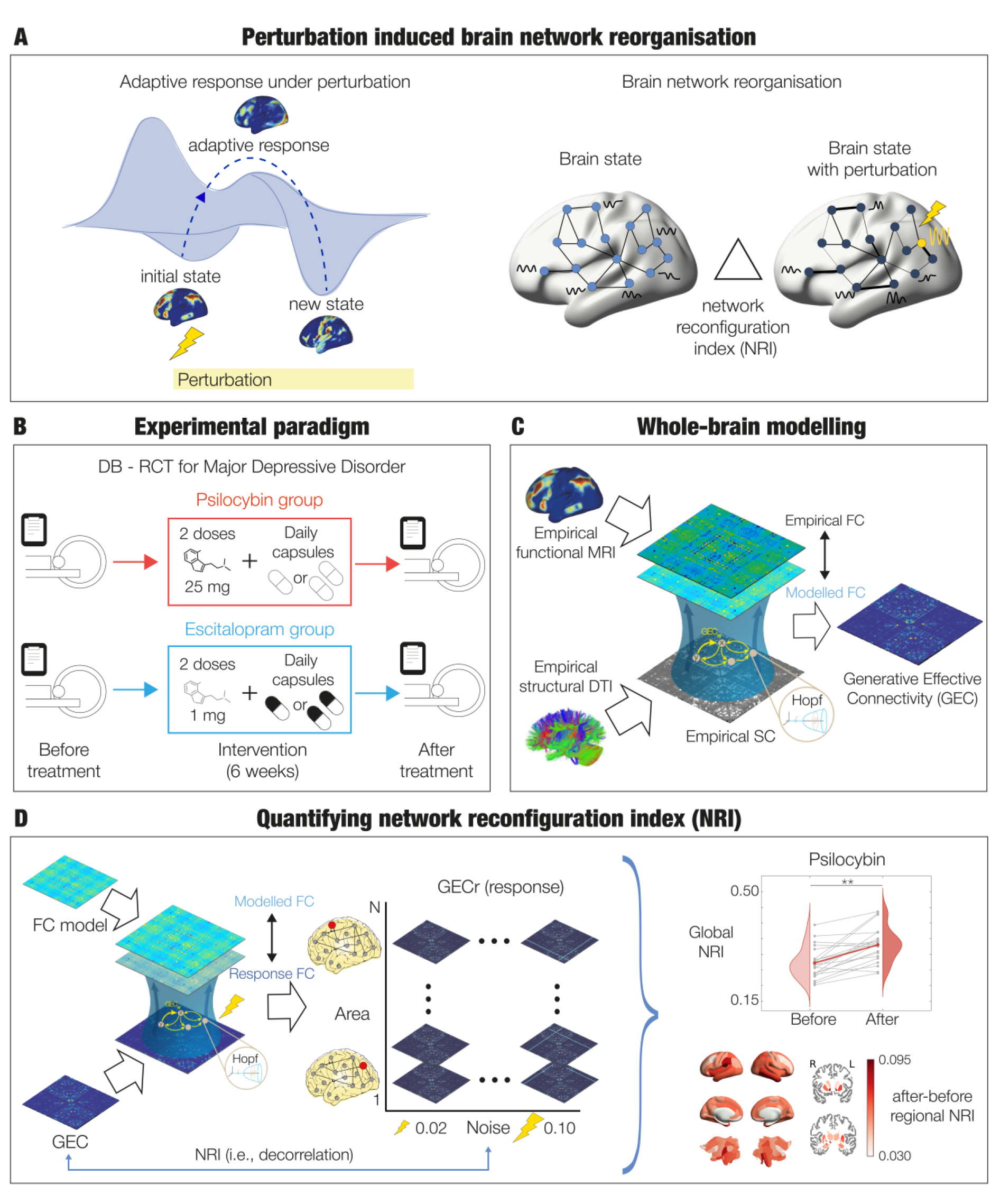
Brain network reconfiguration index (NRI). **A**. The left schematic shows the evolution of an initial brain state, its adaptive response to perturbation, and a new perturbed brain state. The right schematic shows an initial brain network and a reorganised brain network containing the perturbation. The network reconfiguration index (NRI) quantifies the extent of brain reorganisation by calculating the dissimilarity between the initial and the perturbed brain network. **B**. Experimental paradigm consisting of a double-blind randomised control trial comparing interventions of depression, namely psilocybin and escitalopram. **C**. We built Hopf whole-brain models for each individual in each group (psilocybin and escitalopram) and session (before and after treatment). We combined anatomical structural connectivity (SC) and functional connectivity (FC) to create a generative effective connectivity (GEC) matrix, optimised and fitted to empirical fMRI. **D**. We characterised NRI as the ability of the whole-brain to reorganise with a local perturbation. First, we perturbed each brain area independently with different stimulation intensities, starting off with the initial GEC and re-optimised the matrix by fitting it to the initial model, giving as output a response GEC (GECr). We quantified the dissimilarity between the GECr and the initial GEC by computing the decorrelation between the matrices as a measure of NRI. The higher the decorrelation, the higher the NRI, and the further away the response model to the initial model.

### Statistical Analysis

All statistical analysis was performed using permutation-based Wilcoxon sign-rank with 1000 permutations and a significance threshold of 0.05, paired and non-paired when applicable. We applied a False Discovery Rate (FDR) method to correct for multiple comparisons (Hochberg & Benjamini, 1990).

## Results

### Global brain network reconfiguration index following each treatment

The NRI of individuals, averaged across all brain areas and stimulation intensities, increased following psilocybin and decreased following escitalopram (**Figure 2A**). This effect was consistent across stimulation intensities (**Figure 2B**), with between sessions showing significant differences surviving correction for multiple comparisons at all intensities above 0.02. An intensity of 0.02 is very close to the initial model (i.e., 0.01) and produced negligible perturbations.

**Figure 2.**
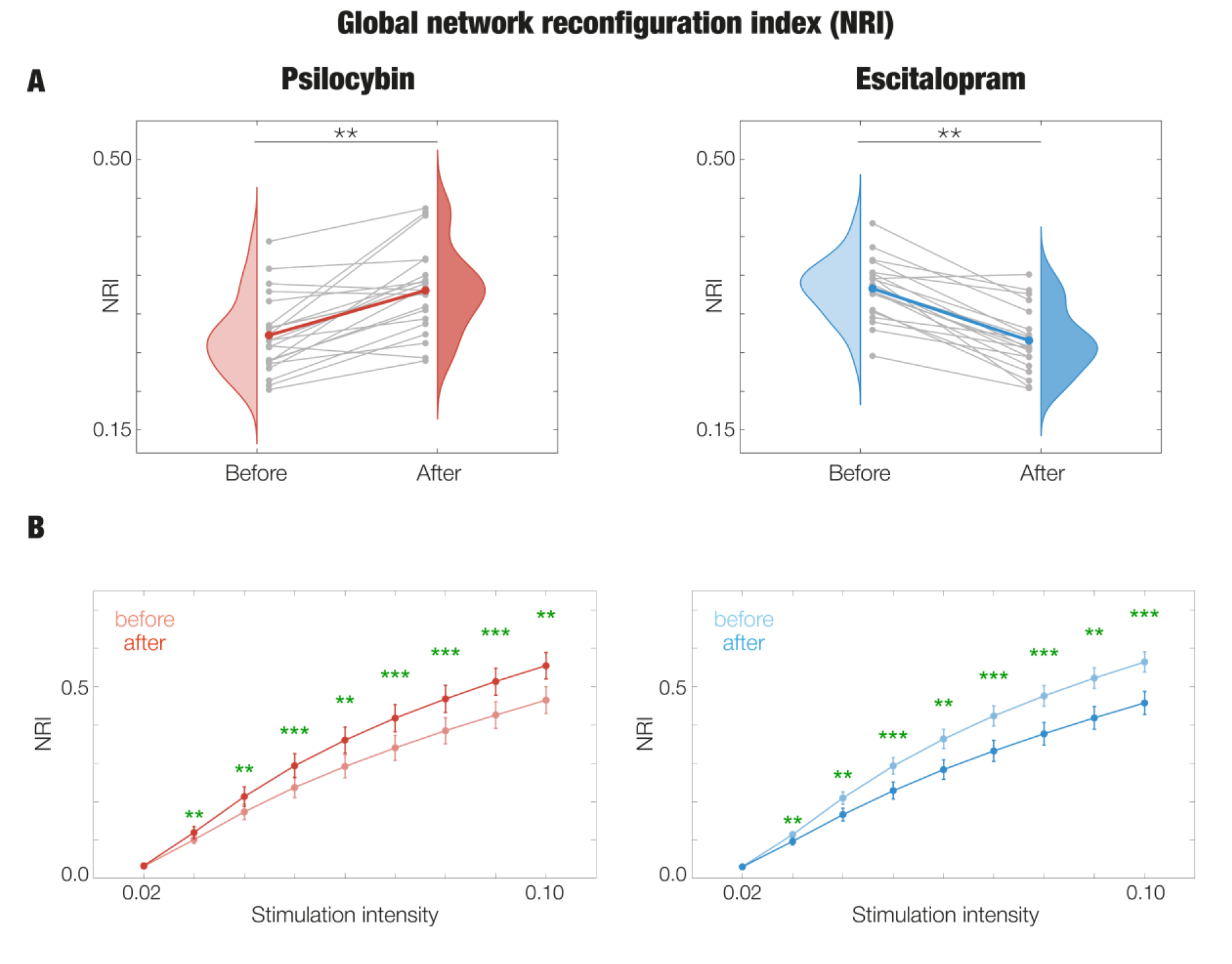
Whole-brain NRI. **A**. The NRI of individuals, averaged across all brain areas and stimulation intensities, for each intervention arm and session. Grey lines represent the trajectory of each individual whereas the coloured line represents their averaged trajectory within each intervention arm (red for psilocybin and blue for escitalopram). **B**. The NRI of individuals at each stimulation intensity averaged across all brain areas, for each intervention arm and session. Lineplots represent the NRI averaged across participants and errorbars their standard deviation. Asterisks represent significant differences (**, p < 0.01; ***, p < 0.001), in green the ones surviving correction by multiple comparisons.

### Regional brain network reconfiguration index following each treatment

Furthermore, the NRI for all brain areas, averaged across noise levels, was higher after treatment in the psilocybin arm and lower for escitalopram (**Figure 3A**). In both treatment groups, strongest absolute changes in NRI were found mostly in subcortical areas and some areas with highest association with the ventral attention network (VAN) and limbic network (LIM) (Yeo et al., 2011). Moreover, in the psilocybin group regions associated with the default mode network (DMN) also showed prominent changes, whereas in the escitalopram group additional changes were observed in areas contributing mainly to the dorsal attention network (DAN).

**Figure 3.**
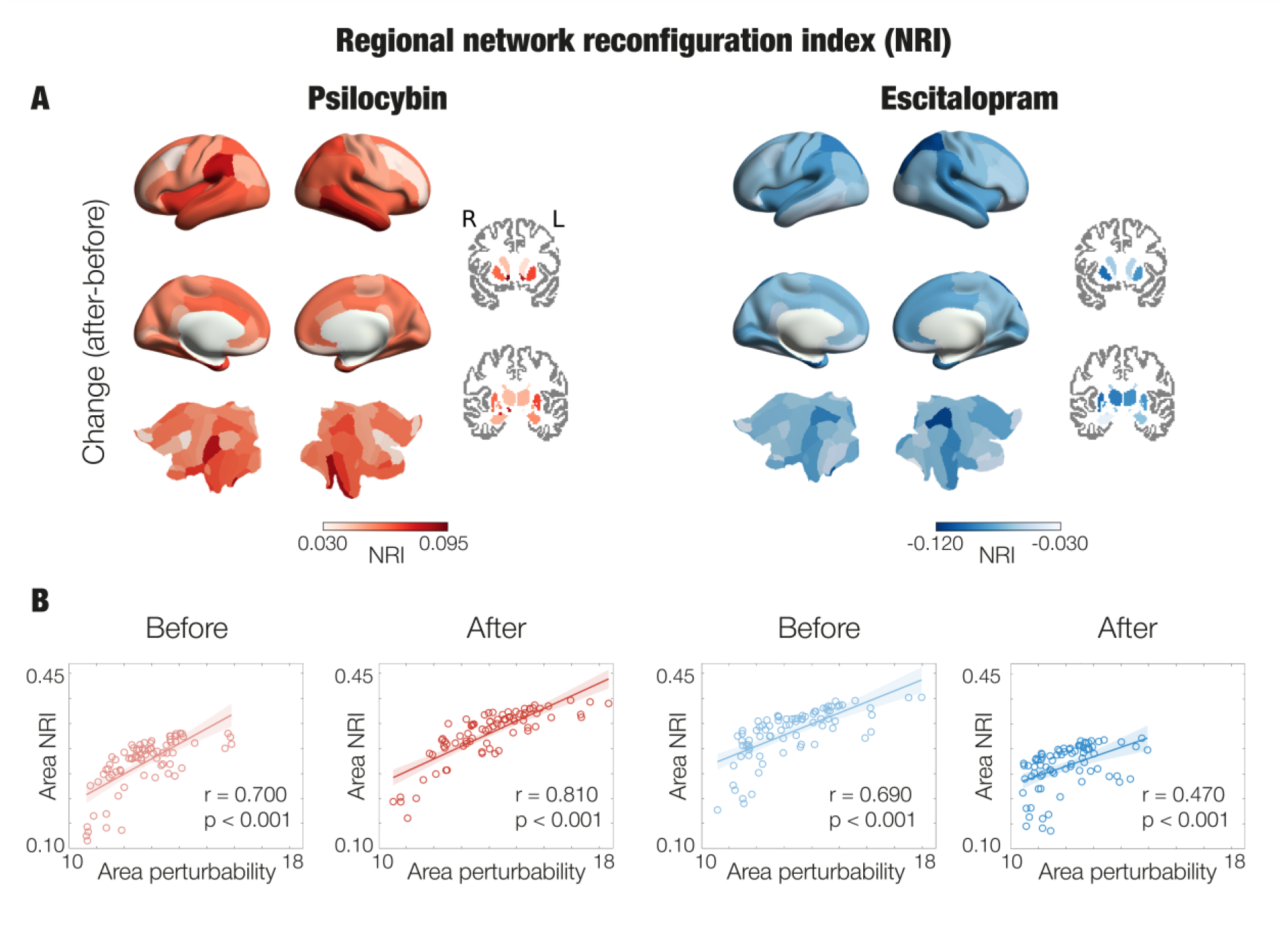
Regional NRI. **A.** The NRI change (after-before) (mean of participants) after perturbing each brain area averaged across stimulation intensities. The brain renders reveal all areas increase in psilocybin (left) and decrease in escitalopram (right). Subcortical areas are shown on slices in Montreal Neurological Institute (MNI) space (coronal axis y= 12 and −10 mm for top and bottom images, respectively). **B**. For each group and session, we obtained the NRI after perturbing each brain area averaged across participants and stimulation intensities. Then, we correlated regional NRI with the perturbability of each brain area averaged across participants from (Dagnino et al., 2026).

To investigate the origin behind regional differences in NRI, we examined the relation between the NRI of brain areas and their perturbability. Regional NRI values were obtained by averaging across noise levels and participants, while perturbability measures were derived from (Dagnino et al., 2026), averaged across participants. Perturbability reflects the degree of non-equilibrium dynamics, such that areas with greater perturbability orchestrate the brain’s hierarchical organisation. Further methodological details are provided in (Deco et al., 2023). The Pearson correlation between nodal perturbability and NRI was significantly positive for both groups (psilocybin and escitalopram) and sessions (before and after treatment) (**Figure 3B**).

### Traditional approaches

We then evaluated complementary methods to our measure by implementing the traditional approaches of (Cascone et al., 2021) and (Kotlarz, 2024) that quantify brain network integrity under targeted node removal. These approaches describe their measures as brain network resilience. Given the term resilience encompasses a broad range of definitions, which is in turn different from mental resilience, throughout the analysis we refer directly to the specific graph-theory metric computed in each analysis. Their pipelines consist in removing nodes (i.e., brain areas) from the thresholded and/or binarised functional connectivity (FC) matrix in an accumulated manner, following a predefined order, and measuring the outcome FC with measures from graph theory (Rubinov & Sporns, 2010). We applied the pipelines in the GEC after thresholding the matrices in a range of 5-100% top edges, with and without binarisation (**Figure 4**). For more details on the methodological pipelines, please see **Supplementary Information**.

**Figure 4.**
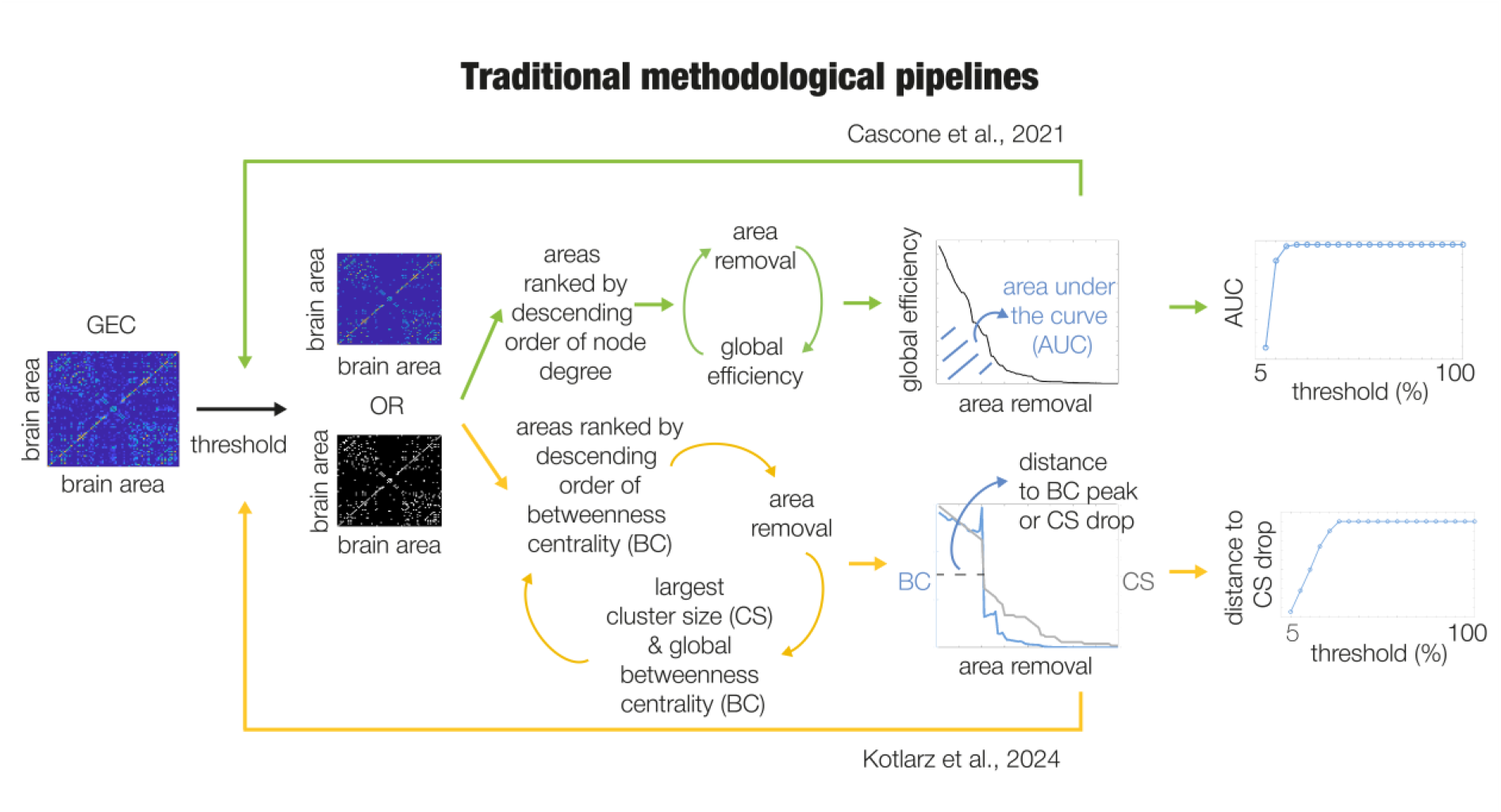
Traditional approaches. We thresholded the GEC matrices of each individual and then moved forward with the weighted version or binarising the resulting matrix. For any case (weighted or binarised GEC) we implemented iterative processes for different threshold ranges. Following (Cascone et al., 2021), we computed the following. First, we calculated the node degree and ordered the brain areas in descending order. Then, we iterated across all brain areas and removed one by one, calculating the resulting global efficiency (GE) of the matrix – inverse of shortest path length. Finally, we computed the area under the curve (AUC) for the GE vs. node removed. Following (Kotlarz, 2024), we performed a three-step process. First, ordered brain areas in descending order of betweenness centrality (BC) – a measure quantifying how often a brain area appears on the shortest path of other areas. Then, removed the top node, and computed the largest cluster size (CS) and global betweenness centrality (BC). We repeated this three-step procedure in an accumulated manner until all brain areas were removed (i.e., converted to zero) resulting in a plot of BC and CS vs. node removal. A peak in BC, preceding a drop in CS, is expected due to sudden shifts in network integration after a certain number of nodes are removed. We then computed the distance (number of brain areas to remove) to the drop in largest cluster size.

In both methods, changes were opposite for the two interventions studied (**Figure 5**). In (Cascone et al., 2021), the pipeline consists of measuring the area under the curve (AUC) of global efficiency (GE) across nodal removal. The retained topological integrity (i.e., AUC) is interpreted as a characteristic of the healthy brain. In the weighted case, psilocybin showed a significant reduction in AUC after treatment for all thresholds, and escitalopram showed a significant increase in AUC for all thresholds. As such, lower/higher AUC for the psilocybin/escitalopram group correspond to weaker/stronger paths (i.e., strength of connections) after treatment. In the binarised case, both groups showed the same aforementioned significant differences in thresholds 5-15%. For higher thresholds, the changes were reversed, psilocybin had a higher significant AUC after treatment for thresholds equal and above 30%, and escitalopram had a lower significant AUC after treatment for thresholds above 30%. The higher/lower AUC for the psilocybin/escitalopram group corresponds to increased/decreased connectivity patterns (i.e., communication via any path) after treatment when most edges are considered (>30% thresholds), whereas the opposite occurs when only top edges are considered.

**Figure 5.**
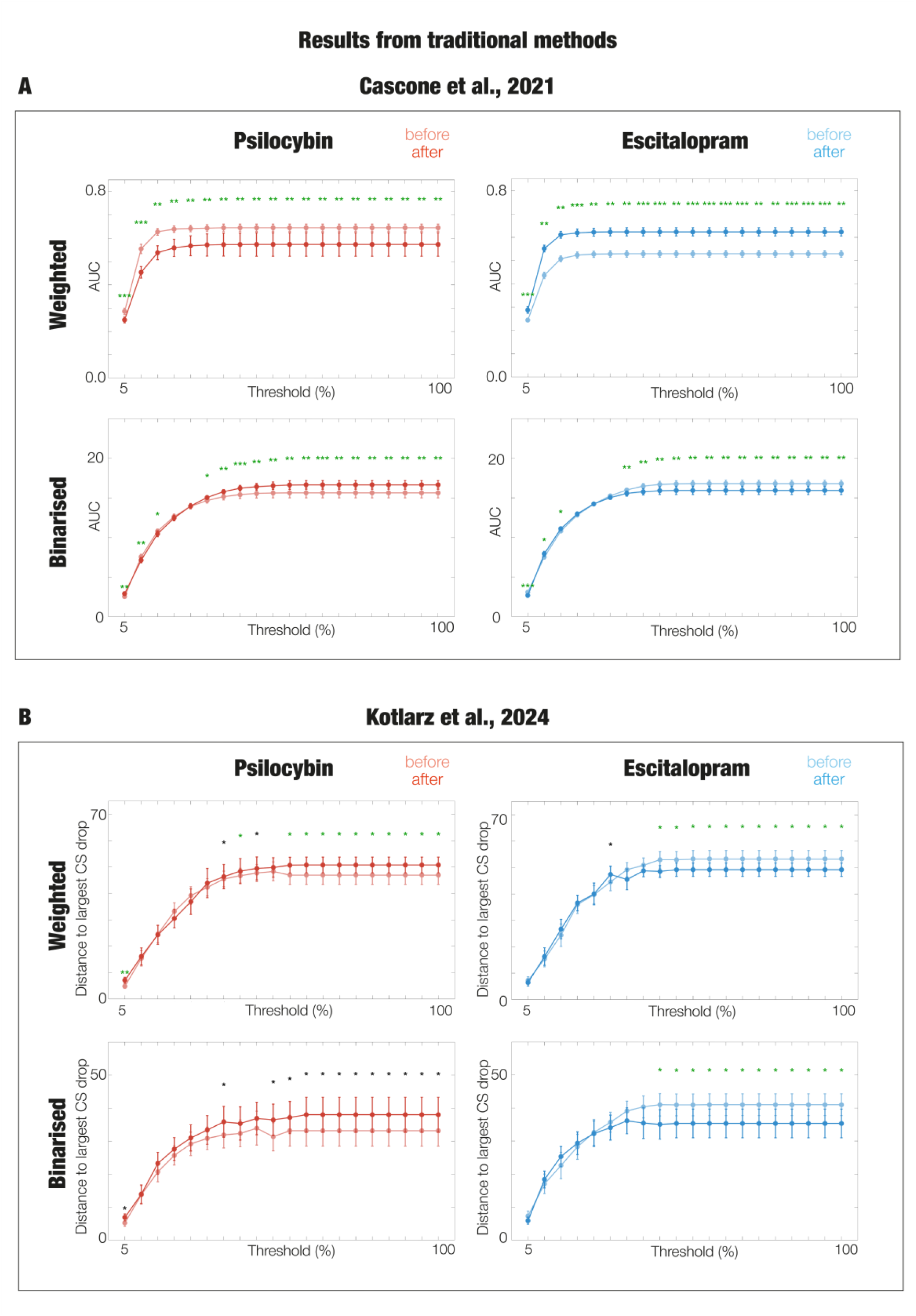
Results from traditional methods. Different thresholds for psilocybin (left column) and escitalopram (right column) in both the binarised and weighted cases for **A**. (Cascone et al., 2021) showing the area under the curve, and for **B**. (Kotlarz, 2024) showing the distance to largest drop in cluster size (CS). Lineplots show the mean across participants and bars their standard deviation. Significance is represented by asterisks (*** p < 0.001; ** p < 0.01 and * p < 0.05, green asterisks correspond to the ones surviving correction by multiple comparisons).

In (Kotlarz, 2024), the pipeline consists of computing the distance (i.e., number of nodes removed) to the highest peak of betweenness centrality (BC), or largest drop in cluster size (CS), after all nodes are removed. This distance is associated with a characteristic of near-critical dynamics in health from the viewpoint of network compartmentalisation, maintaining modular structure and operating near the edge of breakdown. In both the weighted and binarised cases, psilocybin showed a higher distance to drop in CS after treatment compared to baseline for almost all thresholds (for an example see **Supplementary Figure S1**), and escitalopram showed the opposite change, both significant for most of thresholds in the range of 35-50%, with psilocybin also showing a significance for threshold of 5%.

## Discussion

We successfully investigated whether whole-brain organisation under perturbation in patients with MDD changes following two different interventions: psilocybin versus escitalopram. To address this, we built whole-brain models, applied perturbations, and built new models containing the perturbations. We then quantified the brain network reconfiguration index (NRI), a measure of the degree of brain reorganisation under perturbation (**Figure 1)**. Our results showed that NRI increases following psilocybin and decreases following escitalopram. These opposing neural effects are consistent with previous analyses in the same dataset (Dagnino et al., 2026; Deco, 2024), further supporting the idea that clinical improvement across treatments may be achieved through distinct neural processes. Together, our findings suggest that brain NRI captures a complementary dimension of treatment effects in whole-brain dynamics in MDD.

We found increased NRI following psilocybin and decreased NRI following escitalopram across regions and stimulation intensities (**Figures 2 and 3A**). Increased NRI following psilocybin suggests a shift from rigid maladaptive brain network configurations towards greater capacity for brain network reorganisation under sustained perturbation, indicating that brain dynamics may depart easier from their unperturbed organisation. We extend previous results showing higher susceptibility following psilocybin (Socoro-Garrigosa et al., 2025), therefore the brain is not only more sensitive to a given perturbation but also undergoes increased reorganisation under such perturbation. This is consistent with the association between psychedelics and heightened brain flexibility and malleability (Vohryzek et al., 2022). In contrast, decreased NRI following escitalopram suggests a stabilisation of brain dynamics closer to the unperturbed configuration, extending previous results on reduced susceptibility following escitalopram (Socoro-Garrigosa et al., 2025).

The increased NRI following psilocybin could align with theoretical models of psychedelic therapy proposing that its effects are driven by increased brain and mind plasticity, understood as the quality of being easily shaped (i.e., changeability). Within this framework, heightened plasticity is hypothesised to transiently increase sensitivity to contextual influences, enabling the reset or recalibration of maladaptive patterns of thought and behaviour when combined with appropriate psychotherapy (Juliani et al., 2024; Kocarova et al., 2021). The latter is key, given therapeutic alliance has been associated with greater emotional breakthrough and improved clinical outcomes (Murphy et al., 2021). This is supported by evidence showing increased psychological flexibility (i.e., ability to remain open and present, and act aligned to one’s values and goals) and cognitive flexibility (i.e., ability to shift attention between different aspects) following psychedelic therapy (Kocarova et al., 2021). At the mechanistic level, this heightened plasticity may be understood within the RElaxed Beliefs Under Psychedelics (REBUS) model and the anarchic brain hypothesis (Carhart-Harris & Friston, 2019). This framework proposes that the acute psychedelic phase leads to a flattening of the brain’s energy landscape, enabling the system to escape rigid or maladaptive attractors, and increased sensitivity to perturbations. Overall, increased NRI following psilocybin may reflect enhanced brain changeability beyond the acute phase, facilitating adaptive reorganisation under appropriate contextual conditions.

Interestingly, distinct changes in each treatment have been also observed at a psychological level, with psilocybin associated with increased affective responses, and escitalopram with reduced emotional responsiveness. Specifically, in this same dataset, psilocybin showed increases in emotional intensity and responsiveness, whereas escitalopram showed decreases (Harding et al., 2025; Wall et al., 2025). Furthermore, psilocybin showed greater reduction in avoidance and anhedonia relative to escitalopram (Carhart-Harris et al., 2021). Qualitative reports from another study of psilocybin for treatment resistant depression further support this distinction, with patients perceiving psilocybin as facilitating a shift from disconnection to connection, and from avoidance to acceptance, while their previous antidepressant treatment was associated with reduced distress through pain suppression (Watts, 2017). Antidepressants have been identified to reduce negative biases early in treatment, preceding mood improvement, which in turn may be accompanied with decreased responsiveness to both positive and negative surprising events or emotionally salient cues (i.e., emotional blunting) (Godlewska & Harmer, 2021).

We also found that regional NRI was positively correlated with perturbability values from (Dagnino et al., 2026) across sessions and intervention arms (**Figure 3B**). Regions with high perturbability are thought to play a role in orchestrating global hierarchical brain dynamics (Deco et al., 2023). As such, our results suggest that the brain undergoes greater network reconfiguration when the region containing the perturbation has greater orchestrating influence. Consistent with this interpretation, previous work in healthy individuals showed that core areas, identified as having strongly weighted connections with the rest of the brain, exhibit lower responses to local perturbations while eliciting stronger responses in the non-stimulated areas (Ponce-Alvarez, 2025).

Our work could be understood as a way of assessing the capacity to persist, resist, recover, and/or reorganise under disturbance, inherent and fundamental to biological systems (Artime, 2024; Liu, 2022). In the brain, traditional approaches have studied this by removing brain areas or connections based on structural (Crossley et al., 2014; Xu et al., 2021) and functional information (Argiris et al., 2024; Cascone et al., 2021; Menardi et al., 2021; van Assche et al., 2022). In these studies, node removal generally follows a given order (e.g., descending order of node degree), and together with the measurement of the effects of node removal, both are computed using graph theory measures (Rubinov & Sporns, 2010). We extended our analysis to assess topological integrity of brain networks after targeted node removal following (Cascone et al., 2021) and (Kotlarz, 2024) (**Figures 4 and 5**). After targeted node removal, (Cascone et al., 2021) measures global efficiency, and (Kotlarz, 2024) measures number of regions to be removed before a drop in network compartmentalization. Following (Cascone et al., 2021), our results showed for psilocybin after treatment a lower global efficiency in the weighted case, revealing weaker connections, and a higher global efficiency in the binarised case when considering most of the top edges, corresponding to increased communication paths. The opposite occurred for escitalopram. Moreover, following (Kotlarz, 2024), our results showed for psilocybin after treatment an increased number of nodes needed to be removed until a drop in largest cluster size, in both the weighted and binarised cases, suggesting a higher integrated network. The opposite occurred for escitalopram. Overall, these results suggest that following psilocybin treatment, network organisation is characterised by a greater number of connections, which are in turn weaker, as indicated by the approach of (Cascone et al., 2021), together with more distributed communication pathways, as indicated by the approach of (Kotlarz, 2024). These findings are consistent with our results showing a brain state that is more easily reconfigured under perturbation (i.e., increased NRI). The opposite pattern is observed following escitalopram across all measures.

Our approach has several advantages. First, whole-brain Hopf models have shown to effectively replicate empirical fMRI whole-brain dynamics with a reasonable balance between complexity of the models and realism of the simulations (Deco et al., 2017). After perturbation, the strength of each existing connection in the GEC is updated in an iterative manner to achieve the best fit to the initial model. This way, we do not measure the response to perturbation as in classical computational neuroscience approaches, but the brain’s reconfiguration under a perturbation, seeking to retain a state that approximates the initial simulated configuration. Furthermore, by building a whole-brain model that captures the “arrow of time” from empirical data, NRI is quantified within a thermodynamic framework (Kringelbach et al., 2023). This allows us to study the relation between NRI and functional hierarchies from (Dagnino et al., 2026). Lastly, traditional methods assessing brain integrity after node removal are based on different rules (e.g., descending order of the betweenness centrality) whereas for computing NRI there is no need to bias the ordering of node perturbation. Overall, our framework offers more nuanced insights for measuring effects of whole-brain perturbations and network reconfiguration in treatments for depression.

Our approach has several limitations worth mentioning. First, the analysis was applied to a relatively small dataset, limiting the statistical power of the findings. Future research using larger cohorts will be important for increasing the robustness and generalisability of the results. In addition, the choice of brain parcellation may affect estimates of the NRI, and future analyses should examine whether the reported effects replicate across atlases with different spatial resolutions. Lastly, as in previous work, the structural connectivity matrix used to build the initial whole-brain models was obtained from healthy participants. Although this limitation is partially addressed during the optimisation process, future studies could further improve the models using individualised structural connectivity data from each patient.

Overall, our work successfully addressed the question on whether brain reorganisation under perturbation changes with two different interventions for MDD. By using whole-brain models and artificial perturbations we quantified the network reconfiguration index. Our results showed that NRI increases with psilocybin and decreases with escitalopram, aligning with results of traditional methods using graph theory. Overall, our work contributes to the growing body of knowledge on brain perturbation, depression, and treatment effects.

## Data availability

The raw data can be requested to R.L.C.-H., the chief investigator on the original work.

## Code availability

Analysis on whole-brain models was performed in MATLAB R2022a and R2024a software from MathWorks (Natick, MA, USA). We used functions from the free available Brain Connectivity Toolbox (brain-connectivity-toolbox.net). Subcortical images were built in Python V3.12. The code is available from the corresponding authors upon request.

## Acknowledgements

P.D. is supported by the Department of Research and Universities of the Government of Catalonia, Agency for Management of University and Research Grants (AGAUR), FI-SDUR programme (Grant No. 2022 FISDU 00229). I.A.P is supported by Grant PID2022-136216NB-100 funded by MICIU/AEI/ 10.13039/501100011033 and, as appropriate, by “ERDF A way of making Europe”, by “ERDF/EU”, by the “European Union” or by the “European Union NextGenerationEU/PRTR”. M.L.K. is supported by the Centre for Eudaimonia and Human Flourishing (funded by the Pettit and Carlsberg Foundations) and Center for Music in the Brain (funded by the Danish National Research Foundation, DNRF117). Y.S.P. is supported by the EU funded Project NEurological MEchanismS of Injury, and Sleep-like cellular dynamics (NEMESIS; ref. 101071900) funded by the EU ERC Synergy Horizon Europe, and by the Grant PID2024-162576NA-I00 funded by MICIU/AEI/10.13039/501100011033 and by “ERDF A way of making Europe”, ERDF, EU. G.D. is supported by Grant PID2022-136216NB-I00 funded by MICIU/AEI/10.13039/501100011033 and by “ERDF A way of making Europe”, “ERDF, EU”, Project NEurological MEchanismS of Injury, and Sleep-like cellular dynamics (NEMESIS) (ref. 101071900) funded by the EU ERC Synergy Horizon Europe, AGAUR research support grant (ref. 2021 SGR 00917) funded by the Department of Research and Universities of the Generalitat of Catalunya, and Grant PID2024-155136NI-I00 financed by MICIU/AEI/10.13039/501100011033/ and by “ERDF A way of making Europe”, ERDF, EU. The double-blind randomized controlled trial was funded by a private donation from the Alexander Mosley Charitable Trust, supplemented by Founders of Imperial College London’s Centre for Psychedelic Research. The research was conducted at The National Institute for Health and Care Research (NIHR) Imperial Clinical Research Facility (ICRF). We would like to thank the Clinical Data Systems team at the Imperial Clinical Trials Unit for their support. All authors affiliated with the Imperial College London Division of Psychiatry are supported by the National Institute of Health Research (NIHR) Imperial Biomedical Research Collaboration.

## Competing interests

R.C.-H. Robin is scientific advisor to Entropy Neurodynamics, Red Light Holland, Otsuka, and AtaiBeckley. D.E. is acting as a paid scientific advisor for Aya Biosciences, Lophora Aps, Clerkenwell Health, Mindstate Design Lab, and Otsuka. The other authors declare no competing interests.

## References

Acero-Pousa, I., Escrichs, A., Clara Dagnino, P., Sanz Perl, Y., Kringelbach, M. L., Uhlhaas, P. J., & Deco, G. (2025). Reconfiguration of functional brain hierarchy in schizophrenia. Transl Psychiatry, 15(1), 356. 10.1038/s41398-025-03584-0

Acero-Pousa, I., Sanz Perl, Y., Vohryzek, J., Garcia Guzmán, E., Escrichs, A., Mindlin, I., Diego Sitt, J., Kringelbach, M. L., & Deco, G. (2025). Inception: Simulating Personalized Long-Term Recovery in Disorders of Consciousness using Whole-Brain Computational Perturbations. bioRxiv. 10.1101/2025.07.23.666344

Argiris, G., Stern, Y., & Habeck, C. (2024). Brain Resilience to Targeted Attack of Resting BOLD Networks as a Measure of Cognitive Reserve. Res Sq. 10.21203/rs.3.rs-5356022/v1

Artime, O. G., Marco De Domenico, Manlio Gleeson, James P.; Makse, Hernán A.; Mangioni, Giuseppe Perc, Matjaž; Radicchi Filippo. (2024). Robustness and resilience of complex networks. Nature Reviews Physics, 6, 114–131. 10.1038/s42254-023-00616-2

Breyton, M., Fousek, J., Rabuffo, G., Sorrentino, P., Kusch, L., Massimini, M., Petkoski, S., & Jirsa, V. (2025). Spatiotemporal brain complexity quantifies consciousness outside of perturbation paradigms. Elife, 13. 10.7554/eLife.98920

Carhart-Harris, R., Giribaldi, B., Watts, R., Baker-Jones, M., Murphy-Beiner, A., Murphy, R., Martell, J., Blemings, A., Erritzoe, D., & Nutt, D. J. (2021). Trial of Psilocybin versus Escitalopram for Depression. N Engl J Med, 384(15), 1402–1411. 10.1056/NEJMoa2032994

Carhart-Harris, R. L., & Friston, K. J. (2019). REBUS and the Anarchic Brain: Toward a Unified Model of the Brain Action of Psychedelics. Pharmacol Rev, 71(3), 316–344. 10.1124/pr.118.017160

Cascone, A. D., Langella, S., Sklerov, M., & Dayan, E. (2021). Frontoparietal network resilience is associated with protection against cognitive decline in Parkinson‘s disease. Commun Biol, 4(1), 1021. 10.1038/s42003-021-02478-3

Crossley, N. A., Mechelli, A., Scott, J., Carletti, F., Fox, P. T., McGuire, P., & Bullmore, E. T. (2014). The hubs of the human connectome are generally implicated in the anatomy of brain disorders. Brain, 137(Pt 8), 2382–2395. 10.1093/brain/awu132

Dagnino, P. C., Acero-Pousa, I., Zamora-López, G., Escrichs, A., Erritzoe, D., Nutt, D. J., Carhart-Harris, R. L., Sanz Perl, Y., Kringelbach, M. L., & Deco, G. (2026). Distinct brain responses to psilocybin and escitalopram in depression captured by the Fluctuation-Dissipation Theorem. bioRxiv, 2026.2006.2012.731811. 10.64898/2026.06.12.731811

Deco, G., Cabral, J., Saenger, V. M., Boly, M., Tagliazucchi, E., Laufs, H., Van Someren, E., Jobst, B., Stevner, A., & Kringelbach, M. L. (2018). Perturbation of whole-brain dynamics in silico reveals mechanistic differences between brain states. NeuroImage, 169, 46–56. 10.1016/j.neuroimage.2017.12.009

Deco, G., Cruzat, J., Cabral, J., Tagliazucchi, E., Laufs, H., Logothetis, N. K., & Kringelbach, M. L. (2019). Awakening: Predicting external stimulation to force transitions between different brain states. Proc Natl Acad Sci U S A, 116(36), 18088–18097. 10.1073/pnas.1905534116

Deco, G., Kringelbach, M. L., Jirsa, V. K., & Ritter, P. (2017). The dynamics of resting fluctuations in the brain: metastability and its dynamical cortical core. Sci. Rep., 7(1), 3095. 10.1038/s41598-017-03073-5

Deco, G., Lynn, C. W., Sanz Perl, Y., & Kringelbach, M. L. (2023). Violations of the fluctuation-dissipation theorem reveal distinct nonequilibrium dynamics of brain states. Phys. Rev. E., 108(6). 10.1103/physreve.108.064410

Deco, G. S. P., Yonatan Johnson, Samuel; Bourke, Niamh; Carhart-Harris Robin L., Kringelbach Morten L. (2024). Different hierarchical reconfigurations in the brain by psilocybin and escitalopram for depression. Nature Mental Health 2, 1096–1110.10.1038/s44220-024-00298-y

Deco, G. V. D.; Kringelbach, M. L. (2021). Revisiting the global workspace orchestrating the hierarchical organization of the human brain. Nat. Hum. Behav., 5(4), 497–511. 10.1038/s41562-020-01003-6

Desikan, R. S., Ségonne, F., Fischl, B., Quinn, B. T., Dickerson, B. C., Blacker, D., Buckner, R. L., Dale, A. M., Maguire, R. P., Hyman, B. T., Albert, M. S., & Killiany, R. J. (2006). An automated labeling system for subdividing the human cerebral cortex on MRI scans into gyral based regions of interest. NeuroImage, 31(3), 968–980. 10.1016/j.neuroimage.2006.01.021

GBD. (2018). Global, regional, and national incidence, prevalence, and years lived with disability for 354 diseases and injuries for 195 countries and territories, 1990–2017: a systematic analysis for the Global Burden of Disease Study 2017. The Lancet, 392(10159), 1789–1858. 10.1016/S0140-6736(18)32279-7

Godlewska, B. R., & Harmer, C. J. (2021). Cognitive neuropsychological theory of antidepressant action: a modern-day approach to depression and its treatment. Psychopharmacology (Berl), 238(5), 1265–1278. 10.1007/s00213-019-05448-0

Gu, S., Betzel, R. F., Mattar, M. G., Cieslak, M., Delio, P. R., Grafton, S. T., Pasqualetti, F., & Bassett, D. S. (2017). Optimal trajectories of brain state transitions. NeuroImage, 148, 305–317. 10.1016/j.neuroimage.2017.01.003

Harding, R., Singer, N., Wall, M. B., Hendler, T., Erritzoe, D., Nutt, D., Carhart-Harris, R., & Roseman, L. (2025). Dissociable effects of psilocybin and escitalopram for depression on processing of musical surprises. Mol Psychiatry, 30(7), 3188–3196. 10.1038/s41380-025-03035-8

Hochberg, Y., & Benjamini, Y. (1990). More powerful procedures for multiple significance testing. Stat. Med., 9(7), 811–818. 10.1002/sim.4780090710

Holtzheimer, P. E., & Mayberg, H. S. (2011). Stuck in a rut: rethinking depression and its treatment. Trends Neurosci, 34(1), 1–9. 10.1016/j.tins.2010.10.004

Juliani, A., Safron, A., & Kanai, R. (2024). Deep CANALs: a deep learning approach to refining the canalization theory of psychopathology. Neurosci Conscious, 2024(1), niae005. 10.1093/nc/niae005

Klein, A., & Tourville, J. (2012). 101 labeled brain images and a consistent human cortical labeling protocol. Front Neurosci, 6, 171. 10.3389/fnins.2012.00171

Kocarova, R., Horacek, J., & Carhart-Harris, R. (2021). Does Psychedelic Therapy Have a Transdiagnostic Action and Prophylactic Potential? Front Psychiatry, 12, 661233. 10.3389/fpsyt.2021.661233

Kotlarz, P. F., Marcelo; Nino, Juan C.; Alzheimer’s Disease Neuroimaging Initiative. (2024). Brain Network Modularity and Resilience Signaled by Betweenness Centrality Percolation Spiking. Applied Sciences, 14(10). 10.3390/app14104197

Kringelbach, M. L., Perl, Y. S., Tagliazucchi, E., & Deco, G. (2023). Toward naturalistic neuroscience: Mechanisms underlying the flattening of brain hierarchy in movie-watching compared to rest and task. Sci Adv, 9(2), eade6049. 10.1126/sciadv.ade6049

Kringelbach, M. L., Sanz Perl, Y., & Deco, G. (2024). The Thermodynamics of Mind. Trends Cogn Sci, 28(6), 568–581. 10.1016/j.tics.2024.03.009

Liu, X. L., Daqing Ma, Manqing; Szymanski Boleslaw K.; Stanley, H. Eugene; Gao, Jianxi. (2022). Network resilience. Physics Reports, 971, 1–108. 10.1016/j.physrep.2022.05.001

Lynn, C. W., Cornblath, E. J., Papadopoulos, L., Bertolero, M. A., & Bassett, D. S. (2021). Broken detailed balance and entropy production in the human brain. Proc. Natl. Acad. Sci. U. S. A., 118(47). 10.1073/pnas.2109889118

Menardi, A., Reineberg, A. E., Vallesi, A., Friedman, N. P., Banich, M. T., & Santarnecchi, E. (2021). Heritability of brain resilience to perturbation in humans. NeuroImage, 235, 118013. 10.1016/j.neuroimage.2021.118013

Murphy, R., Kettner, H., Zeifman, R., Giribaldi, B., Kartner, L., Martell, J., Read, T., Murphy-Beiner, A., Baker-Jones, M., Nutt, D., Erritzoe, D., Watts, R., & Carhart-Harris, R. (2021). Therapeutic Alliance and Rapport Modulate Responses to Psilocybin Assisted Therapy for Depression. Front Pharmacol, 12, 788155. 10.3389/fphar.2021.788155

Ponce-Alvarez, A. (2025). Network Mechanisms Underlying the Regional Diversity of Variance and Time Scales of the Brain‘s Spontaneous Activity Fluctuations. J Neurosci, 45(10). 10.1523/JNEUROSCI.1699-24.2024

Rubinov, M., & Sporns, O. (2010). Complex network measures of brain connectivity: uses and interpretations. NeuroImage, 52(3), 1059–1069. 10.1016/j.neuroimage.2009.10.003

Sanz Perl, Y., Escrichs, A., Tagliazucchi, E., Kringelbach, M. L., & Deco, G. (2022). Strength-dependent perturbation of whole-brain model working in different regimes reveals the role of fluctuations in brain dynamics. PLoS Comput Biol, 18(11), e1010662. 10.1371/journal.pcbi.1010662

Sanz Perl, Y., Fittipaldi, S., Gonzalez Campo, C., Moguilner, S., Cruzat, J., Fraile-Vazquez, M. E., Herzog, R., Kringelbach, M. L., Deco, G., Prado, P., Ibanez, A., & Tagliazucchi, E. (2023). Model-based whole-brain perturbational landscape of neurodegenerative diseases. Elife, 12. 10.7554/eLife.83970

Sanz Perl, Y., Pallavicini, C., Piccinini, J., Demertzi, A., Bonhomme, V., Martial, C., Panda, R., Alnagger, N., Annen, J., Gosseries, O., Ibanez, A., Laufs, H., Sitt, J. D., Jirsa, V. K., Kringelbach, M. L., Laureys, S., Deco, G., & Tagliazucchi, E. (2023). Low-dimensional organization of global brain states of reduced consciousness. Cell Rep, 42(5), 112491. 10.1016/j.celrep.2023.112491

Socoro-Garrigosa, M., Perl, Y. S., Kringelbach, M. L., Erritzoe, D., Nutt, D. J., Carhart-Harris, R., Vohryzek, J., & Deco, G. (2025). Perturbing whole-brain models of brain hierarchy: An application for depression following pharmacological treatment. Ann N Y Acad Sci, 1550(1), 255–272. 10.1111/nyas.15391

van Assche, M., Klug, J., Dirren, E., Richiardi, J., & Carrera, E. (2022). Preparing for a Second Attack: A Lesion Simulation Study on Network Resilience After Stroke. Stroke, 53(6), 2038–2047. 10.1161/STROKEAHA.121.037372

Vohryzek, J., Cabral, J., Lord, L. D., Fernandes, H. M., Roseman, L., Nutt, D. J., Carhart-Harris, R. L., Deco, G., & Kringelbach, M. L. (2024). Brain dynamics predictive of response to psilocybin for treatment-resistant depression. Brain Commun, 6(2), fcae049. 10.1093/braincomms/fcae049

Vohryzek, J., Cabral, J., Vuust, P., Deco, G., & Kringelbach, M. L. (2022). Understanding brain states across spacetime informed by whole-brain modelling. Philos Trans A Math Phys Eng Sci, 380(2227), 20210247. 10.1098/rsta.2021.0247

Wall, M. B., Demetriou, L., Giribaldi, B., Roseman, L., Ertl, N., Erritzoe, D., Nutt, D. J., & Carhart-Harris, R. L. (2025). Reduced Brain Responsiveness to Emotional Stimuli With Escitalopram But Not Psilocybin Therapy for Depression. Am J Psychiatry, 182(6), 569–582. 10.1176/appi.ajp.20230751

Watts, R. D., Camilla Krzanowski, Jacob; Nutt, David; Carhart-Harris, Robin. (2017). Patients’ Accounts of Increased “Connectedness” and “Acceptance” After Psilocybin for Treatment-Resistant Depression. Sage Journals, 57(5). 10.1177/0022167817709585

Xu, D., Xu, G., Zhao, Z., Sublette, M. E., Miller, J. M., & Mann, J. J. (2021). Diffusion tensor imaging brain structural clustering patterns in major depressive disorder. Hum Brain Mapp, 42(15), 5023–5036. 10.1002/hbm.25597

Yeo, B. T., Krienen, F. M., Sepulcre, J., Sabuncu, M. R., Lashkari, D., Hollinshead, M., Roffman, J. L., Smoller, J. W., Zollei, L., Polimeni, J. R., Fischl, B., Liu, H., & Buckner, R. L. (2011). The organization of the human cerebral cortex estimated by intrinsic functional connectivity. J Neurophysiol, 106(3), 1125–1165. 10.1152/jn.00338.2011

Zheng, Y., Yang, Y., Zhen, Y., Wang, X., Liu, L., Zheng, H., & Tang, S. (2025). Understanding Altered Dynamics in Cocaine Use Disorder Through State Transitions Mediated by Artificial Perturbations. Brain Sci, 15(3). 10.3390/brainsci15030263

