## Supplementary Information for "Divergent changes in perturbation-induced brain reconfiguration following depression treatment with psilocybin and escitalopram"

### Materials and Methods

#### - Participants

For participants to be eligible, they needed a confirmed diagnosis of unipolar MDD from a general practitioner. This consisted of a score of 16 or higher on the 21-item Hamilton Depression Rating scale. Individuals were queried about previous psychedelics use, with positive reports of 31% in the psilocybin group and 24% in the escitalopram group. Exclusion criteria consisted of immediate family or personal history of psychosis, risky physical health condition assessed by a physician, serious suicide attempts history, positive pregnancy test, contraindications for MRI, contraindications for selective serotonin reuptake inhibitors (SSRIs), previous use of escitalopram. Treatment resistance was not an inclusion or exclusion criteria. Eligible participants underwent telephone interview screening and comprehensive evaluations of mental and physical medical history.

#### - Treatment protocol

A total of 59 patients with MDD were recruited. A random number generator assigned 30 individuals to the psilocybin group and 29 individuals to the escitalopram group. The final number of participants for this analysis were n=22 (mean age= 41.9 years, s.d.= 11.0, 14 male and 8 female), and n=20 (mean age= 38.7 years, s.d. =11.0, 14 male and 6 female) for the psilocybin and escitalopram arms, respectively. In the psilocybin group the following patients were excluded: one for choosing not to take daily capsules (i.e., placebo) and COVID-19 UK lockdown, two for not attending post-treatment sessions, and five for excessive fMRI head motion. In the escitalopram group the following patients were excluded: four discontinued because of escitalopram adverse reactions, one reported use of cannabis, one was lost in COVID-19 UK lockdown, three were excluded for excessive fMRI head motion.

#### - Magnetic resonance imaging acquisition

Brain imaging was carried out with a 3T Siemens Tim Trio set-up at Invicro. Acquisition of brain anatomy was obtained using the recommended MPRAGE parameters from Alzheimer's Disease Neuroimaging Initiative, Grand Opportunity (ADNI-GO56): isotropic voxels 1-mm; sagittal slices 160; in-plane field of view 256 x 256; echo time (TE) 2.98 ms; repetition time (TR) 2,300 ms; generalized autocalibrating partially parallel acquisitions (GRAPPA) acceleration 2; flip angle 9°; bandwidth per pixel 240 Hz. Acquisition of brain functional data (i.e., fMRI) was collected during eyes-closed resting-state using T2*-weighted echo-planar images and a 32-channel head coil with a total of 480 volumes in ~10 min. Parameters were as follows: isotropic voxels 3-mm; axial slices 44; TE 30 ms; TR 1,250 ms; GRAPPA acceleration 2; flip angle 70°; bandwidth per pixel 2,232 Hz.

#### - BOLD fMRI pre-processing

Imaging data was pre-processed with an in-house pipeline using FMRIB Software Library (FSL) (Smith et al., 2004), Analysis of Functional NeuroImages (AFNI) (Cox, 1996), Freesurfer (Dale et al., 1999) and Advanced Normalization Tools (Avants; Brian B.; Johnson, 2009) packages. Details can be found in (Daws et al., 2022). The pipeline consists of: de-spiking, slice time correction, motion correction, brain extraction, rigid body registration to anatomical scans, nonlinear template registration, scrubbing, band-pass filtering, regression (with six realignment motion regressors, three tissue signal regressors, draining veins and local white matter). Systematic bias from movement artifacts were ruled out by (Daws et al., 2022).

- **Whole-brain computational model**

***Hopf bifurcation model***

We implemented a Stuart-Landau oscillator to describe the local dynamics of each brain area. This corresponds to the normal form (i.e., minimal canonical form) of a supercritical Hopf bifurcation (i.e., standard model to observe transition from noisy to oscillatory dynamics) (Kuznetsov, 2023). Whole-brain dynamics are expressed by coupling the local dynamics of N brain areas (i.e., Stuart-Landau oscillators) with a connectivity matrix *C*. This is defined mathematically by:

$$\frac{dz_j}{dt} = (a_j + i\omega_j)z_j - |z_j|^2 z_j + \sum_{k=1}^{N} C_{jk}(z_k - z_j) + \eta_j, \qquad (1)$$

where for area $j$, $z_j$ is the complex variable specifying the state ($z_j = x_j + iy_j$) and $w_j$ corresponds to the frequency. This frequency is computed after band-pass filtering the empirical fMRI signals in the range of 0.008-0.08Hz, and averaging the peak frequency across individuals. Furthermore, $\eta_j$ is Gaussian noise with variance $\sigma^2$. Lastly, $a_j$ is the bifurcation parameter which governs the dynamics of each local node. For, $a_j > 0$, the dynamics settle into a stable limit cycle, generating self-sustained oscillations of frequency $f_j = w_j/2\pi$. Contrarily, for $a_j < 0$, the dynamics represent a stable spiral point, producing noisy oscillations if noise is present, and damped oscillations otherwise. The fMRI signals are modeled by the real part of the state variable ($xj$ = Real($zj$)).

***Linearization of the model***

We used a bifurcation parameter value just below zero ($a_j = -0.02$) for all brain areas given it was found to be the best working point for fitting whole-brain models (i.e., generates fluctuating stochastically structured signals preserving and resting-state network structure and dynamically responding brain networks) (Sanz Perl et al., 2023). This allows for a linearization of the dynamics which in turn permits an analytical solution for the functional connectivity matrix $FC$ (i.e., Pearson correlations between all pairs of brain regions) (Ponce-Alvarez & Deco, 2024). By implementing a linear noise approximation and the dynamical system in **Equation 1** the following vector form expresses whole-brain functional correlations:

$$\frac{dz}{dt} = (a - S + i\omega) \odot z - (z \odot \bar{z})z + Cz + \eta. \qquad (2)$$

Here, $[z_1, \ldots, z_N]^T$, $a = [a_1, \ldots, a_N]^T$, $\omega = [\omega_1, \ldots, \omega_N]^T$, $\eta = [\eta_1, \ldots, \eta_N]^T$ and $S = [S_1, \ldots, S_N]^T$, a vector indicating the connectivity strength of each node (i.e., $S_i = \sum_j C_{ij}$). In addition, $[\,]^T$ is the transpose operation, $\odot$ is the symbol for the element-wise product, and $\bar{z}$ is the complex conjugate of $z$.

This equation describes the solution of $\frac{dz}{dt} = 0$, the linear fluctuations around the fixed point $z = 0$. After separating the real and imaginary parts of the state variables and discarding higher-order terms (i.e., terms $(z \odot \bar{z})z$), the evolution of the linear fluctuations follow a Langevin stochastic linear equation as:

$$\frac{d}{dt}\delta u = J\delta u + \eta. \tag{3}$$

The 2N-dimensional vector $\delta u = [\delta x, \delta y]^{\mathrm{T}} = [\delta x_1, \ldots, \delta x_N, \delta y_1, \ldots, \delta y_N]^{\mathrm{T}}$ contains fluctuations of the real and imaginary state variables. Moreover, $J$ is a $2N\,X\,2N$ Jacobian matrix of the system evaluated at the fixed point. It has the following form as block matrix:

$$J = \begin{bmatrix} Jxx & Jxy \\ Jyx & Jyy \end{bmatrix}. \tag{4}$$

Here, $J_{xx}$ , $J_{xy}$, $J_{yx}$ and $J_{yy}$ are matrices of size $N\,X\,N$. In particular, $J_{xx} = J_{yy} = diag(a - S) + C$ and $J_{xy} = -J_{yx} = diag(\omega)$ . The $diag(v)$ corresponds to the diagonal matrix with the vector $v$ as the diagonal. The linearization is possible only if $z = 0$ is a stable solution of the system, corresponding to $J$ having all eigenvalues with a negative real part.

Lastly, in order to compute the covariance matrix $K = \langle \delta u \delta u^T \rangle$, **Equation 3** is expressed as $d\delta u = J\delta u dt + dW$. Here, $dW$ is a 2N-dimensional Wiener process with covariance $\langle dW dW^T \rangle = Qdt$. In addition, $Q$ is the noise covariance matrix, which is diagonal if the noise is uncorrelated. Using Itô's stochastic calculus, $d(\delta u \delta u^T) = d(\delta u)\delta u^T + \delta u d(\delta u^T) + d(\delta u)d(\delta u^T)$. And, taking expectations, noting $\langle \delta u dW^T \rangle = 0$, and keeping terms to first order in the differential $dt$, we get:

$$z = \frac{dK}{dt} = JK + KJ^T + Q. \tag{5}$$

The stationary covariance can be obtained analytically using the eigen-decomposition of the Jacobian matrix $J$ by solving this equation for the case $\frac{dK}{dt} = 0$ (Deco et al., 2023).

***Model optimisation: Generative Effective Connectivity***

In order to build a whole-brain model of each participant fitted best to its corresponding empirical data, we implemented a pseudo-gradient procedure to optimise the coupling connectivity matrix C. Our starting point was the standard structural connectivity matrix (SC), computed with diffusion MRI data of the Human Connectome Project ( HCP) (Deco et al., 2021). We iteratively updated these initial anatomical connections to create a resulting matrix called generative effective connectivity (GEC), consisting of effective conductivity values for each pair of existing connections. In each iteration, we compared the simulated with empirical measures of functional connectivity and normalised time-shifted covariances as follows:

$$C_{ij} = C_{ij} + \alpha(FC_{ij}^{emp} - FC_{ij}^{sim}) + \varsigma(FS_{ij}^{emp}(\tau) - FS_{ij}^{sim}(\tau)). \tag{6}$$

Here, $\alpha$ and $\varsigma$ have a value of 0.0004 and 0.0001, respectively. The empirical functional connectivity ($FC^{emp}$) corresponds to the normalised covariance matrix of the fMRI timeseries. Furthermore, the empirical normalised time-shifted covariance ($FS^{emp}(\tau)$) is obtained by computing the shifted covariance matrix ($KS^{emp}(\tau)$) and dividing each pair of regions *(i,j)* by $\sqrt{KS_{ii}^{emp}(0)KS_{jj}^{emp}(0)}$. With respect to the simulated measures, the simulated functional connectivity ($FC^{sim}$) is obtained from the first $N$ rows and columns of the simulated covariance matrix $K$. Lastly, the simulated normalised time-shifted covariance ($FS^{sim}(\tau)$) is derived by selecting the first N rows and columns of the simulated time-shifted covariance ($KS^{sim}(\tau)$), and dividing each pair of regions ($i$,$j$) by $\sqrt{KS_{ii}^{sim}(0)KS_{jj}^{sim}(0)}$. Here,

$$KS^{sim}(\tau) = exp(\tau\, J)K, \tag{7}$$

where $KS^{sim}(0) = K$. The time-shift (set to $\tau = 2$) from the time-shifted covariances break the symmetry in the couplings, which have shown to improve the fitting (Gilson et al., 2018) and introduce non-equilibrium dynamics (Kringelbach et al., 2023).

The process updates known existing connections within each hemisphere following the anatomical connections. Homotopic areas (i.e., areas in opposite hemispheres) are also updated given that it has been revealed that they are captured less accurately in tractography. We performed 1500 iterations and every 100 iterations, if the error between the simulated and empirical measures was higher than the previous iteration, or smaller than 0.001 from the last iteration, the procedure broke. The final optimised matrix, GEC, corresponds to the last iteration. The whole pipeline was performed in a two-step manner, by first calculating the GEC at a group level, with the DTI as a starting point. And then, we calculated the GEC of each participant, using the GEC of the corresponding group as the starting point.

**- Brain network reconfiguration index**

***Perturbation protocol***

The *in-silico* stimulation consisted in increasing the Gaussian noise $\eta$ of a Stuart-Landau oscillator. It was done systematically and single-sited, with a perturbation intensity ranging from 0.02-0.10 in steps of 0.01, in each brain area separately (i.e., N=1-80), whilst maintaining the rest at 0.01.

***Model response***

For each patient, we separately re-optimised the initial GEC for each perturbation intensity at each brain area. This was done by optimizing the GEC containing the perturbation so that it is fitted to the initial simulated measures, creating a response GEC (GECr). The node oscillation frequencies were the same as in the first brain models (i.e., empirical frequencies averaged across individuals of each group).

The optimisation consisted, in each iteration, of comparing the response with the initial simulated measures of functional connectivity and normalised time-shifted covariances as follows:

$$C_{ij} = C_{ij} + \alpha(FC_{ij}^{sim} - FC_{ij}^{sim\_r}) + \varsigma(FS_{ij}^{sim}(\tau) - FS_{ij}^{sim\_r}(\tau)). \tag{8}$$

This time, $\alpha$ and $\varsigma$ have a value of 0.004 and 0.001, respectively. The response measures are calculated the same way as the aforementioned simulated measures, the response functional connectivity ($FC^{sim\_r}$) is obtained from first $N$ rows and columns of the response covariance matrix $K^r$. And, the response normalised time-shifted covariance ($FS^{sim\_r}(\tau)$) is derived by selecting the first N rows and columns of the response time-shifted covariance ($KS^{sim\_r}(\tau)$), and dividing each pair ($i,j$) by $\sqrt{KS_{ii}^{sim\_r}(0)KS_{jj}^{sim\_r}(0)}$, where:

$$KS^{sim\_r}(\tau) = exp(\tau J)K^r, \qquad (9)$$

As in the initial optimisation, the process updates existing anatomical intra-hemispherical connections, as well as interhemispheric connections. Here, 500 iterations were performed, given that the error (distance) and fitting (correlation) between the GECr and GEC converged already to a stable value (the same applies for the response and simulated functional connectivity and normalised time-shifted covariances). We assumed functionality of the reconfigured models with perturbation given they are fitted to the initial GEC during the re-optimisation.

***Computation of network reconfiguration index (NRI)***

We assessed the capacity of the brain to reorganise in front of a constant local increase in noise by quantifying the network reconfiguration index (NRI). As such, we calculated the decorrelation between the GECr and GEC. The higher their decorrelation (i.e., dissimilarity between the initial model and the response model), the higher the NRI (i.e., reorganisation of the model under a perturbation). We obtained one value of NRI for each perturbed brain area and stimulation intensity for each individual.

**- Traditional approaches**

We followed the methodological frameworks from (Cascone et al., 2021) and (Kotlarz, 2024).These implement graph theory measures (Rubinov & Sporns, 2010) to functional connectivity (FC) matrices, whereas for our work we do so in the GEC matrices which are directed, asymmetric, and normalised to 0.02. In both pipelines, matrices are thresholded to maintain the top edges from a total number of possible connections (around 5-30%). In (Cascone et al., 2021) the thresholded matrices are also binarised. In order to have a nuanced representation of the whole range of connections considered we assessed a full range of thresholds, from the maintenance of top edges (5% of top connections) - which might prune important connections -, to preserving the full matrix (100% of connections) - which, if binarised, identifies weak connections equally important as strongest ones. Furthermore, we studied weighted matrices and after binarisation, given that strength levels are important in effective connectivity and it might offer meaningful results rather than computing measures on the patterns alone.

***Method from Cascone et al., 2021***

In (Cascone et al., 2021), they apply as threshold 2.5-20% of top edges in steps of 2.5%. Furthermore, they binarise the matrix. Their motive is to minimise spurious edges which might impact network measures, given that graph metrics on high density graphs have been shown to be more similar to those computed on random graphs, and the fact that low sparsity graphs reveal higher test-retest reproducibility (Sinclair et al., 2015; Wang et al., 2011). Here, we applied as threshold 5-100% of top edges in steps of 5% in order to cover the full range of possibilities. Furthermore, we implemented two alternatives, the thresholded matrix with weighted values, and after binarisation. The following pipeline was applied to each of the resulting matrices. First, the degree of the nodes (i.e., connection of each node to other nodes) is calculated (Rubinov & Sporns, 2010). For the case of the binarised matrices, this is computed as the unweighted degree:

$$k_i = \sum_{j=1}^{n} d_j^i. \tag{10}$$

Here, subindexes *i* and *j* are brain areas and *d* is their connection. Then, nodes are sorted in descending order of degree and removed in an iterative and accumulated manner (i.e., connections changed to zero). Furthermore, for the non-binarised matrices, this is computed as the weighted degree:

$$k_i^W = \sum_{j=1}^{n} w_{ij}, \tag{11}$$

where the connection weight between the brain areas *i* and *j* is $w_{ij}$.

After each node removal in either case, the global efficiency (inverse of shortest path between two nodes) is computed as (Rubinov & Sporns, 2010):

$$E_{global} = \frac{1}{n(n-1)} \sum_{i,j,j \neq 1} \frac{1}{L_{ij}}, \tag{12}$$

where *n* is the total amount of brain areas and $L_{ij}$ is the shortest path length between brain areas *i* and *j*. Lastly, the area under the curve (AUC) of the final curve of global efficiency as a function of nodes removed is computed.

***Method from Kotlarz et al., 2024***

In (Kotlarz, 2024) they apply as threshold 5-30% of top edges, to remove spurious edges and negative correlations, without any posterior binarisation. Here, we applied as threshold 5-100% of top edges in steps of 5%, and to the weighted and binarised matrix, to cover the full range of possibilities. The following pipeline was implemented to each of the resulting matrices. First, the betweenness centrality (BC) of each node (i.e., fraction of shortest paths flowing in each node) is calculated as (Rubinov & Sporns, 2010):

$$b_i = \sum_{h,j \in N\ h \neq j, h \neq i, j \neq i} \frac{\rho_{hj}(i)}{\rho_{hj}} \tag{13}$$

Here, $\rho_{hj}(i)$ corresponds to shortest path between brain areas *h* and *j* which passes through brain area *i*, whereas $\rho_{hj}$ is the number of shortest paths between nodes *h* and *j*. In the weighted case, the shortest path is calculated as the minimum total weight, whereas for the binarised version, it is calculated as the fewest number of steps.

Then, brain areas are sorted in descending order of BC, and the top one removed (i.e., connections changed to zero). Lastly, the largest cluster size (CS) and global BC is computed in the resulting matrix. The global BC corresponds to the average BC of all brain areas. In (Kotlarz, 2024), the largest cluster size is obtained using 'get_components' from the Brain Connectivity Toolbox (Rubinov & Sporns, 2010). Given our matrices are directed, we computed the largest strongly connected component using the 'conncomp' function from Matlab. This three-step procedure (nodal quantifier, nodal removal, and BC and CS measures of the resulting matrix) is repeated in an iterative manner until all nodes are removed. The largest CS and averaged BC is plotted against fraction of attacks and as expected, there are large decreases in CS preceding spikes in average BC. After removing a certain number of brain areas, there is a point in which removing one more node generates a sudden shift and restructuring in the network. In order to obtain a quantitative analysis of these results, we moved a step further from (Kotlarz, 2024) investigation, and calculated the distance to the largest drop in CS (i.e., number of brain areas to remove).

### Figures

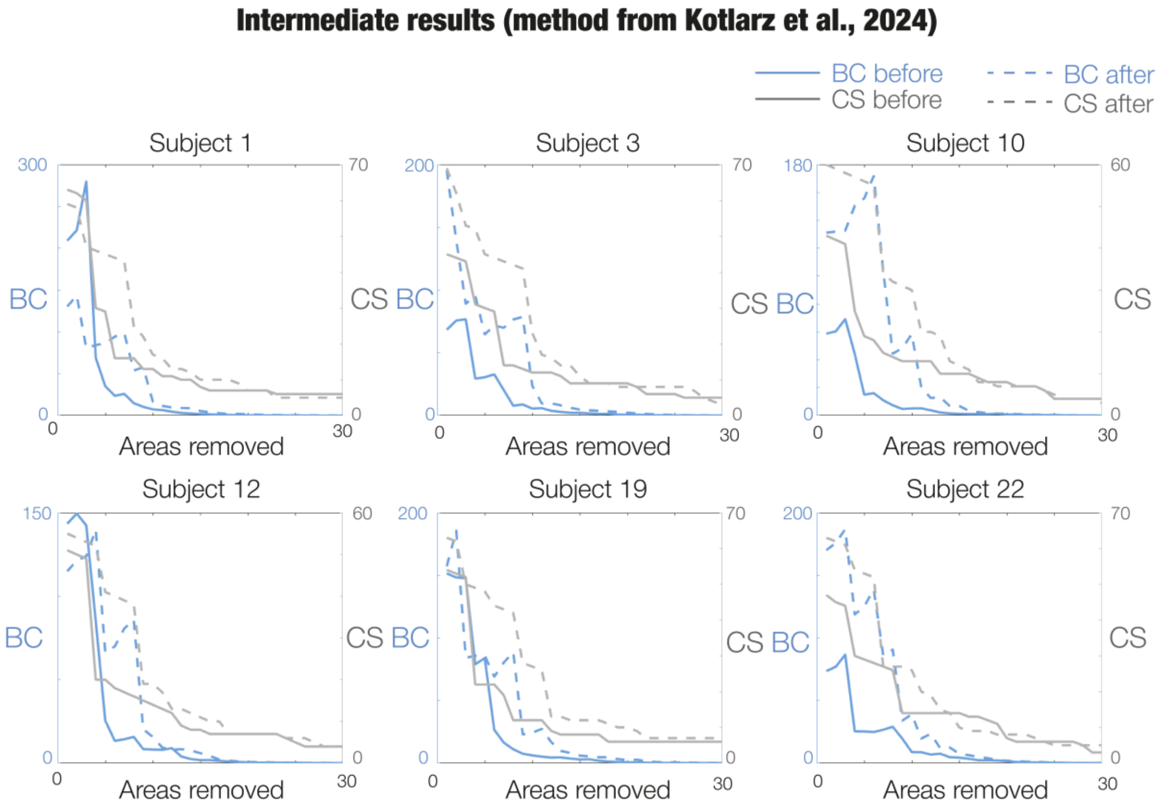


***Supplementary Information Figure S1. Example of traditional method from (Kotlarz, 2024).*** *The plots show global betweenness centrality (BC) – in blue - and largest cluster size (CS) – in grey - vs. node removal for different participants in psilocybin before treatment (5% threshold, binarised case) as example. Lines before treatment are full whereas after treatment are dotted. In all cases, after treatment the distance to the largest drop in CS (or peak BC) is higher.*

### References


Avants; Brian B.; Johnson, H. (2009). Advanced normalization tools (ANTS). *2009*(2), 1–35.

Cascone, A. D., Langella, S., Sklerov, M., & Dayan, E. (2021). Frontoparietal network resilience is associated with protection against cognitive decline in Parkinson's disease. *Commun Biol*, *4*(1), 1021. https://doi.org/10.1038/s42003-021-02478-3

Cox, R. W. (1996). AFNI: software for analysis and visualization of functional magnetic resonance neuroimages. *Comput Biomed Res*, *29*(3), 162–173. https://doi.org/10.1006/cbmr.1996.0014

Dale, A. M., Fischl, B., & Sereno, M. I. (1999). Cortical surface-based analysis. I. Segmentation and surface reconstruction. *NeuroImage*, *9*(2), 179–194. https://doi.org/10.1006/nimg.1998.0395

Daws, R. E., Timmermann, C., Giribaldi, B., Sexton, J. D., Wall, M. B., Erritzoe, D., Roseman, L., Nutt, D., & Carhart-Harris, R. (2022). Increased global integration in the brain after psilocybin therapy for depression. *Nat Med*, *28*(4), 844–851. https://doi.org/10.1038/s41591-022-01744-z

Deco, G., Lynn, C. W., Sanz Perl, Y., & Kringelbach, M. L. (2023). Violations of the fluctuation-dissipation theorem reveal distinct nonequilibrium dynamics of brain states. *Phys. Rev. E.*, *108*(6). https://doi.org/10.1103/physreve.108.064410

Deco, G., Vidaurre, D., & Kringelbach, M. L. (2021). Revisiting the global workspace orchestrating the hierarchical organization of the human brain. *Nat. Hum. Behav.*, *5*(4), 497–511. https://doi.org/10.1038/s41562-020-01003-6

Gilson, M., Deco, G., Friston, K. J., Hagmann, P., Mantini, D., Betti, V., Romani, G. L., & Corbetta, M. (2018). Effective connectivity inferred from fMRI transition dynamics during movie viewing points to a balanced reconfiguration of cortical interactions. *NeuroImage*, *180*(Pt B), 534–546. https://doi.org/10.1016/j.neuroimage.2017.09.061

Kotlarz, P. F., Marcelo; Nino, Juan C.; Alzheimer's Disease Neuroimaging Initiative. (2024). Brain Network Modularity and Resilience Signaled by Betweenness Centrality Percolation Spiking. *Applied Sciences*, *14*(10). https://doi.org/10.3390/app14104197

Kringelbach, M. L., Perl, Y. S., Tagliazucchi, E., & Deco, G. (2023). Toward naturalistic neuroscience: Mechanisms underlying the flattening of brain hierarchy in movie-watching compared to rest and task. *Sci Adv*, *9*(2), eade6049. https://doi.org/10.1126/sciadv.ade6049

Kuznetsov, Y. A. (2023). *Elements of Applied Bifurcation Theory*. Springer. https://doi.org/10.1007/978-3-031-22007-4

Ponce-Alvarez, A., & Deco, G. (2024). The Hopf whole-brain model and its linear approximation. *Sci Rep*, *14*(1), 2615. https://doi.org/10.1038/s41598-024-53105-0

Rubinov, M., & Sporns, O. (2010). Complex network measures of brain connectivity: uses and interpretations. *NeuroImage*, *52*(3), 1059–1069. https://doi.org/10.1016/j.neuroimage.2009.10.003

Sanz Perl, Y., Pallavicini, C., Piccinini, J., Demertzi, A., Bonhomme, V., Martial, C., Panda, R., Alnagger, N., Annen, J., Gosseries, O., Ibanez, A., Laufs, H., Sitt, J. D., Jirsa, V. K., Kringelbach, M. L., Laureys, S., Deco, G., & Tagliazucchi, E. (2023). Low-dimensional organization of global brain states of reduced consciousness. *Cell Rep*, *42*(5), 112491. https://doi.org/10.1016/j.celrep.2023.112491

Sinclair, B., Hansell, N. K., Blokland, G. A., Martin, N. G., Thompson, P. M., Breakspear, M., de Zubicaray, G. I., Wright, M. J., & McMahon, K. L. (2015). Heritability of the network architecture of intrinsic brain functional connectivity. *NeuroImage*, *121*, 243–252. https://doi.org/10.1016/j.neuroimage.2015.07.048

Smith, S. M., Jenkinson, M., Woolrich, M. W., Beckmann, C. F., Behrens, T. E., Johansen-Berg, H., Bannister, P. R., De Luca, M., Drobnjak, I., Flitney, D. E., Niazy, R. K., Saunders, J., Vickers, J., Zhang, Y., De Stefano, N., Brady, J. M., & Matthews, P. M. (2004). Advances in functional and structural MR image analysis and implementation as FSL. *NeuroImage*, *23 Suppl 1*, S208–219. https://doi.org/10.1016/j.neuroimage.2004.07.051

Wang, J. H., Zuo, X. N., Gohel, S., Milham, M. P., Biswal, B. B., & He, Y. (2011). Graph theoretical analysis of functional brain networks: test-retest evaluation on short- and long-term resting-state functional MRI data. *PLoS One*, *6*(7), e21976. https://doi.org/10.1371/journal.pone.0021976
